# Active Surveillance Characterizes Human Intratumoral T Cell Exhaustion

**DOI:** 10.1101/2021.05.06.443033

**Authors:** Ran You, Jordan Artichoker, Adam Fries, Austin Edwards, Alexis Combes, Gabriella Reeder, Bushra Samad, Matthew F. Krummel

## Abstract

Intratumoral T cells that might otherwise control tumors are often identified in an ‘exhausted’ state, defined by specific epigenetic modifications as well as upregulation of genes such as CD38, CTLA-4 and PD-1. While the term might imply inactivity, there has been little study of this state at the phenotypic level in tumors to understand the extent of their incapacitation. Starting with the observation that T cells move more quickly through mouse tumors as residence time increases and they progress towards exhaustion, we elaborated a non-stimulatory live-biopsy method for real-time study of T cell behaviors within individual patient tumors. Using two-photon microscopy, we studied native CD8 T cells interacting with APCs and with cancer cells in different micro-niches of human tumors, finding that T cell speed was variable by region and by patient, was independent of T cell density and was inversely correlated with local tumor density. Across a range of tumor types, we found a strong relationship between CD8 T cell motility and exhausted T cell state that corresponds to observations made in mouse models where exhausted T cells move faster. While this is a small study, it demonstrates at least two types of T cell dynamic states in individual human tumors and supports the existence of an active program in ‘exhausted’ T cells that extends beyond incapacitating them.

## Introduction

Tumors contain inflammatory infiltrates that might detect and eliminate tumor cells, and yet immune tolerance occurs, and cancer evasion persists. T cells isolated from the tumor microenvironment (TME) often exhibit an exhausted phenotype that is characterized by a unique epigenetic landscape and increased inhibitory receptor expression that leads to the reduced proliferative capability and tumor killing (reviewed in (1)). Given that T cells are the targets for most current cancer immunotherapies, understanding how this tolerant state manifests in situ is important. Multiplexed flow cytometry and single-cell transcriptomics have been applied to isolated T cell infiltrates from tumors to investigate T cell stage and function (2), however the associated spatial and dynamic information is lacking. In contrast, fixed imaging methods such as multiplexed ion beam imaging, reveal important links between protein phenotypes, such as PD-1/PD-L1 expression and the spatial organizations of immune infiltrates (3), but lack an understanding of dynamic behaviors and the plasticity of cellular ensembles.

Intravital imaging enables direct visualization of immune cells and their functions in various microcompartments of a tumor and quantitatively measures kinetics of cell behaviors, therefore often revealing previously unknown immune processes (4). Two-photon laser microscopy—using near-infrared excitation to provide improved depth penetration with reduced phototoxicity over longer observation periods—is especially suited for in situ imaging of immune dynamics in intact tumors (5). We and others have established methods to study how immune cells behave in tumors in mice, often using genetically encoded fluorophores (6–8). In one such setting, we found evidence that infiltrated T cells in a spontaneous breast cancer increase their motility following an initial arrest phase (9), suggesting that T cell motility increases in concert with the establishment of exhaustion. T cells that eliminate tumors in mouse models show a variety of behaviors (8), some of which vary during different phases of tumor progression (10, 11). The behaviors of T cells in human tumors—which likely vary in myriad ways relative to mouse models as well as each other—remains largely unstudied.

In one pioneering study using slices of human biopsies, exogenous T cells that were non-native to tumors were labeled with fluorescent dyes and allowed to enter from tissue culture media. There, these newly introduced cells migrated actively in loose collagen areas whereas poorly in dense matrix areas (12). Tracking of these suggested that matrix architecture would guide how such exogenous T cells migrate. Such migration may be akin to that mediated by either chemokines (13) or by forces or structures inherent to the matrix (14). Another study revealed that tumor associated macrophages (TAM) trapped CD8 T cells and excluded them from tumor islets in human lung carcinoma (15). Such studies confirm the frequently long-lasting interaction between T cells and TAM, also observed in mouse tumors (6). However, no systemic approach has yet been developed to study how tumor native T cells migrate nor how that links to their immune function in human tumors. Therefore, we developed and validated a ‘live biopsy’ method for direct imaging of immune function in tumor specimens from cancer patients. We standardized the approach and applied it to tumors from multiple indications to measure immune cell behavior indicated by cell motility and cell-cell interaction among antigen presenting cells (APCs), T cells and tumor cells. The results show both heterogeneity within human cancers with respect to exhaustion but also that the program of exhaustion defines a dynamic state resembling surveillance.

## Results and Discussion

### Develop Live Biopsy Imaging

To characterize the in situ phenotype of intratumoral high-affinity murine T cell, we first generated a cell line from a well-described breast cancer mouse model (mammary tumor virus-polyoma middle T (MMTV-PyMT)) modified to express mCherry and OVA (PyMT-ChOVA) (6) and further adapted it to be suitable for injection into the fat pad of mice with the adoptive transfer of OT-I CD8 T cells at two different time points during tumor progression (Fig. 1A). T cells resident in tumors for 2-week periods (d14), typically adopt a phenotype, dubbed ‘exhausted’ and characterized by upregulation of inhibitory molecules (1, 9, 16), including PD-1, and CD38 (Fig. 1B) (17). A master transcriptional factor, often associated with exhausted T cells, Tox (18–20) is also upregulated when compared to effector T cells that just arrived (d4) in the TME of these tumors. In contrast, the prolonged resident T cells produced less IFNγ (Fig. 1B). Granzyme B production and attenuated Nur77 expression were also observed in long resident T cells in the spontaneous PyMT model as showed in a previous study (9, 21), strongly suggesting the loss of effector functions in these T cells. When we examined the motility of similarly-generated (d14) T cells with the dysfunctional and exhausted phenotype, we found that they fail to arrest in the TME and were dynamic compared to recently arrived (d4) effector T cells (Fig. 1C). Specifically, 72.5% of d14 T cells (Exhausted) moved at speed exceeding the mean of the d4 T cells (Arrival) and 44.6% of cells were fast-moving cells (speed > Mean + 1 standard deviation (SD)).

**Figure 1.**
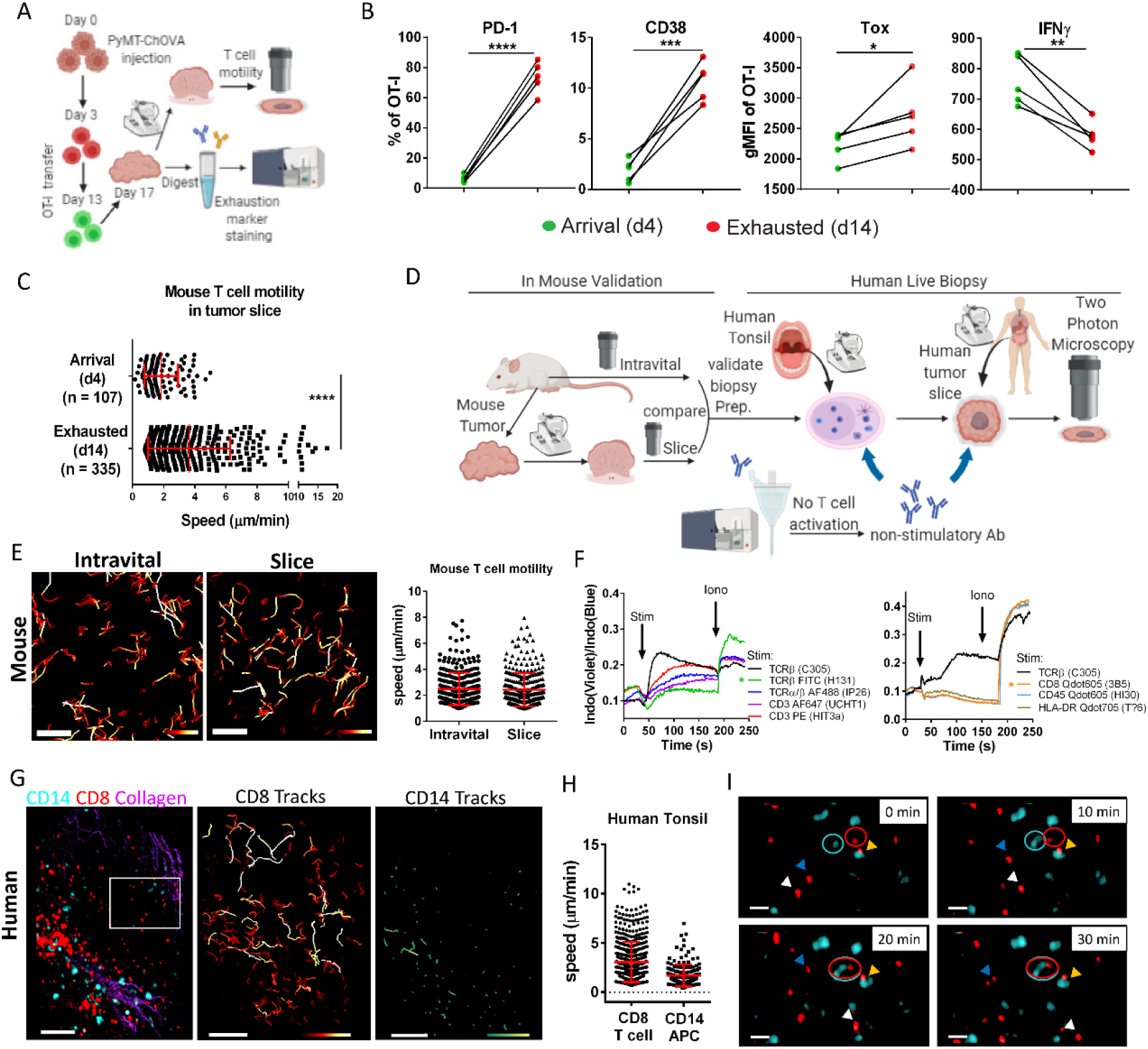
Establishment of live biopsy imaging using non-perturbing antibodies. (A) Schematic diagram of the experimental design: RFP OT-I and GFP OT-1 were separately transferred into the same mice injected with PyMT-ChOVA cells in fat-pad at the time as indicated. Tumors were harvested and split for slice imaging and flow cytometry analysis. (B). Quantitative measurement of exhaustion molecules and IFNγ production by flow cytometry in GFP OT-I T cells or RFP OT-I T cells that were transferred 4 days (arrival) or 14 days (exhausted) into fat-pad injected PyMT-ChOVA tumors before tumor harvest. N = 5 mice. Data is representative of two independent experiments. * P<0.05, ** P<0.01, *** P<0.001, **** P<0.0001, as determined by the Paired Student’s *t*-test. (C) Quantification of T motility of the same groups of cells in tumor slice detected by two photon microscopy. Mean and standard deviation (SD) for the Arrival group were 1.846 and 1.088; Exhausted were 3.637 and 2.639. Data was pooled from four ROIs from two slices per mouse and is representative of two independent experiments. **** P<0.0001, as determined by the Student’s t-test. (D) Schematic diagram for the development of live biopsy imaging strategy. (E) Color-coded tracks of mouse T cells from intravital imaging and slice imaging of fat-pad injected PyMT-ChOVA tumor. Speed range is from 0 to 0.15 μm/sec (red to white). Scale bar is 50 μm. (Left panel); Quantification of T cell speed pooled from two ROIs each group (right panel). (F) Calcium influx assay showed the activation of Jurkat cells, a human T cell line. Violet to blue fluorescence ratio indicates the calcium binding of Indo dye. Antibodies as indicated were added at 30 sec as stimulation reagents (stim) and ionomycin (Iono) were added at 200 sec as positive controls. Stars indicated the non-stimulatory T cell antibodies that used for subsequent imaging experiments. (G) (Left) Representative image and (Right) color-coded track displacement through 30 mins-imaging of a human tonsil slice stained with anti-hCD8-Qdot605 and anti-hCD14-Qdot705. Scale bar is 100 μm. Track length range is from 1 to 50 μm (red to white for T cells; green to yellow for CD14 cells). (H) Track speed mean of CD8 T cell and CD14^+^ cells. (I) Snapshots of the ROI from panel F showing heterogeneous patterns of immune cell behavior. Blue and white arrows: CD8 T cells moved freely. Yellow arrow: CD8 T and CD14^+^ cells dwelled together and formed conjugates for the entire imaging time. Red and blue circle: CD8 T cell and CD14^+^ cells moved towards each other and started to interact. Scale bar is 20 μm. Also see supplementary Video 1.

We sought to extend this result to human tumors and in doing so also shed light on the variety of T cell behaviors within the more diverse domain of human tumors. For this, we needed a method for real time imaging of human tumor biopsies that does not require the addition of new cells for tracking and that could ultimately label and image human cells in biopsy in a way that did not alter their biology (Fig. 1D). To do this, we first validated imaging strategies for tumor biopsy slices, again taking advantage of mice whereby we could compare the data obtained from sliced tumors against data taken from intravital imaging of the same tumor (Fig. 1D). For the data shown in Fig. 1E, we injected the PyMT-ChOVA cell line that was described previously in Fig.1A into the fat pad of the CD2-DsRed mice and varied storage, transport, slicing and imaging conditions to find conditions under which endogenous T cell behaviors were intact. We chose T cell motility rate as our primary measurement since it has previously been shown to be exquisitely sensitive to both temperature and oxygenation (22) but also tracked dendritic cell morphology and behavior under the optimized conditions (data not shown). After iterating, we found an optimum for storage/transport using pre-warmed and oxygenated media and times post-surgery of less than 5 hours (see Methods), and for imaging using a heated perfusion chamber with a constant flow of oxygen-saturated media. Under these conditions, T cell motility rates were well maintained in the tumor slices, being indistinguishable from those observed using intravital imaging (Fig. 1E). We found these conditions were similar for a collection of other mouse tumors including B78 melanoma and MC38 colon carcinoma (Supplemental Fig. 1A).

Since transgenic fluorescent markers are not experimentally tractable in human samples, we next screened and identified a panel of non-stimulatory antibodies that label human immune cells to aid visualization and do not trigger well-established signaling pathways downstream of their ligands (Fig. 1D). For example, we selected a single clone that bound to a TCRβ (Vβ13.1; clone H131) and a single anti-CD8α antibody that bound to human CD8 T cells but did not induce a detectable calcium influx in Jurkat cells (Fig. 1F) or in human peripheral blood mononuclear cells (PBMC) when cross linked (Supplemental Fig. 1B) as compared to the majority of monoclonal antibodies against TCR-associated proteins (e.g. CD3, TCR, etc.) which did so. These clones were not inhibitory, supported by the unchanged calcium influx in response to the stimulatory anti-CD3 when PBMC were pretreated with either of these two antibodies (Supplemental Fig. 1C). The primary screen criterion for antibodies to label myeloid cells was to determine if the expression level of costimulatory molecule or HLA was altered to indicate the effect of the antibodies on the myeloid cell function (Supplemental Fig. 1D). Where possible, Quantum dot (Qdot) conjugates of these clones were used, due to exceptional brightness and photostability (23, 24). As a functional test of this panel *in situ* we examined 400 micron-thick human tonsil slices to visualize CD8 T cells and CD14^+^ myeloid cells (Fig. 1D, G and supplementary video 1). Following imaging, tracking and quantification (Fig. 1H), we found that motility of human T cells fell in the range 1-10μm/min (mean = 3.06), consistent with mouse lymphocyte motility in lymph nodes (25, 26) (27). CD8 T cell speed was also well maintained across 30 μm of depth and throughout a one-hour imaging period (Supplemental Fig. 1E), indicating that there was no obvious adverse effect of imaging on T cell behavior given those parameters. We observed diverse CD8 T cell behaviors in a micro-region of the tonsil, including some with a random walk and others arrested on myeloid cells to form conjugates (Fig. 1I) The fact that T cells arrested and formed contact with myeloid cells might be attributed to the presence of antigens in these inflamed tonsils.

After observing anticipated results in tonsils, we applied this method to tumors obtained from multiple indications (Fig. 1D), including head and neck (HNSC), colorectal (CRC), gynecological (GYN), kidney (KID) and hepatic (HEP) cancers (Supplemental Fig. 1F). Heterogeneous immune infiltrates, stroma represented by collagen, and tumor structure were well preserved.

### Diversity of immune cell dynamics in subregions of human tumors

Live biopsy enabled us to study tumor specimens with a focus on T cell migration and their interaction with other cells in the same microenvironment. Aiming to better understand how endogenous T cell behavior differed in subregions of a human tumor, we first scanned the tumor slice for a broad field of the 3D space to characterize overall immune infiltration and stroma structure. We then ran multiple acquisitions of regions of interest (ROIs) in parallel (28) to capture cellular dynamics in real time. In Supplemental Figure 2, we provide the example of a HNSC tumor, highlighting diverse levels of immune infiltrations among five different regions of interest (ROI), chosen at random during acquisition (Supplemental Fig. 2A, B). From these five ROIs, instantaneous speed varied from less than 1 μm/min to 7 μm/min and mean speed varied from 2.8 μm/min in the fastest region to 1.3 μm/min in the slowest (Supplemental Fig. 2C). When placed in rank order based on speed, neither the density of CD8 T cell nor that of HLA-DR^+^ myeloid cells in these five ROIs were well-correlated with T cell speed (Supplemental Fig. 2C, D, E). In addition, while the contact time between T cells and APCs differed significantly among subregions of a tumor (Supplemental Fig. 2F), this showed no evident relationship to T cell motility.

### CD8 T cell motility was inversely correlated with local tumor density

To extend our analysis to additional tumors and seek out other tissue parameters that might impact motility within a given region of a tumor, we repeated these experiments and now also quantified tumor cell densities. In Figure 2A, using a colorectal tumor as an example, CD8 T cell behaviors in three ROIs with various abundance of EpCAM^+^ tumor cells were quantified (Fig. 2A, B, and supplementary video 2). T cells were tracked, again demonstrating significant regional differences (Fig. 2C, D). As previously observed in the head and neck tumor in Supplemental Figure 2, we found no direct correlation between T cell motility and the density of either CD8 T cells or CD14^+^ myeloid cells, nor the contact time between these two cell types (Supplemental Fig. 3A-C). However, tumor cell density was inversely correlated with T cell motility across the three ROIs (Fig. 2E, F). This relationship repeated in all other tumors examined in this way, for example in another colorectal cancer (Supplemental Fig. 3D-G) and the head and neck cancer shown in Supplemental Figure 2 (Fig. 2F). We note that a similar relationship was observed when labeled exogenous T cells were added to tumor biopsies (12), indicating that this relationship is likely at least partially independent of antigen recognition.

**Figure 2.**
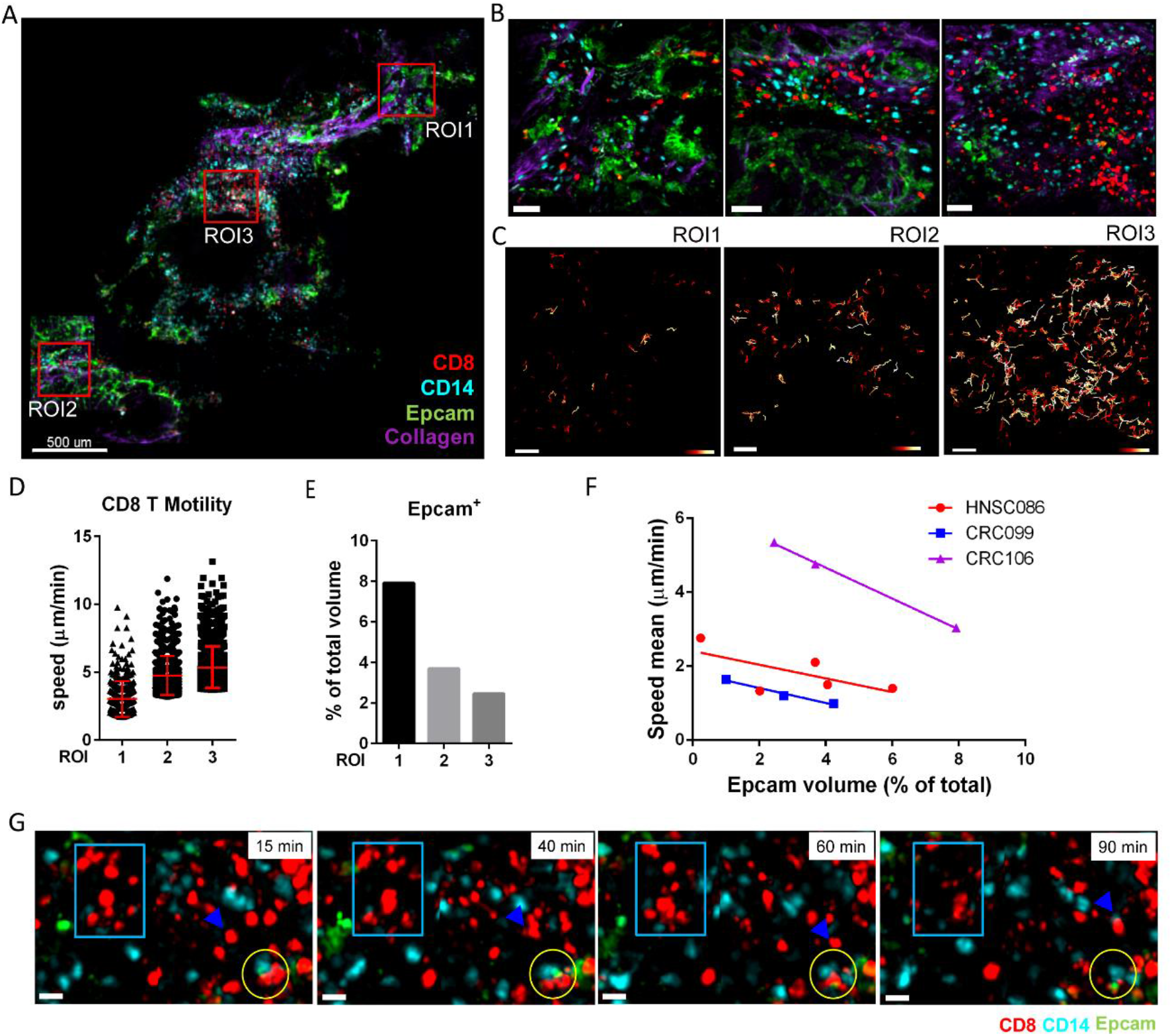
Inverse correlation of CD8 T cell motility with tumor density in cancer biopsies. (A) A scan of a colorectal cancer slice stained with antibodies anti-CD8-Qdot605, anti-CD14-Qdot705, and anti-EpCAM-AF488. (B) Three ROIs of panel A. Scale bar is 50 μm. (C) Color-coded track displacement of CD8 T cells. Length range is from 0.5 (red) to 25 μm (white). Scale bar is 50 μm. (D) The speed of top 10% fastest CD8 T cells from three ROIs was plotted with mean±SD. (E) Percentage of EpCAM^+^ volume in total volume of three ROIs indicating tumor cell density. (F) The speed mean of CD8 T cell from three different tumor samples was plotted respectively against the tumor cell density of the corresponding ROIs from the biopsy. (G) Snapshots of time-lapse video from ROI2 showing: 1) CD8 T cell movement (light blue rectangle: T cells remained in their location and then started to move; blue arrow: T cells moved to interact with CD14^+^ myeloid cells); 2) T-CD14^+^ APC-EpCAM^+^ tumor cell conjugates remained through 90 mins (yellow circle). Also see supplementary Video 2.

Finally, we observed that while most CD8 T cells moved freely when they were not in contact with tumor cells, we occasionally found examples in which they alternatively engaged with myeloid cells and tumor cells to form stable conjugates for at least 90 mins (Fig. 2G and supplemental video 2). Thus, CD8 T cell motility differed among subregions of a tumor, which was potentially caused by heterogeneous local tumor density.

### CD8 T cell motility was correlated with immune states in human tumors

We next sought to broaden this approach toward defining the rate of T cell motility across tumors and across site of origin. In total, 17 fresh tumors from patients with cancers of five different indications were split, with approximately half of the biopsy being dissociated for flow cytometry (Supplemental Fig. 4B) and half being imaged using the live biopsy method (Fig. 3A). Following live biopsy imaging and analysis, human tumors, regardless of their origins, were then ranked, and grouped based on their motility. Though T cell speed was not a binary variable, we classified those with substantial motility across the tumor (motile) from those with very limited T cell motility (Immotile) using a cutoff if the mean was <1 μm per minute (Fig. 3B), in order to investigate if the immune component and cell state were linked to fast versus slow T cell speed. Although these groups had different ischemia time (Supplemental Fig. 4A), which was defined from when the biopsy was resected till it was sliced, we found no statistical support that these were different between motility groups. Average CD8 T cell density was also quantified for these samples, again showing no relationship to motility (Fig. 3C). Based on flow cytometry, we also delineated tumors as immune poor or rich conditions, based on the percentage of CD45^+^ live cells and found no trend or statistical difference in groups (Fig. 3D). The ratio of CD4 to CD8 T cell infiltrates and the abundance of regulatory T cells in the tumors were also measured for each of the tumors that we imaged (Supplemental Fig. 4C-D) and likewise did not correlate with overall motility.

**Figure 3.**
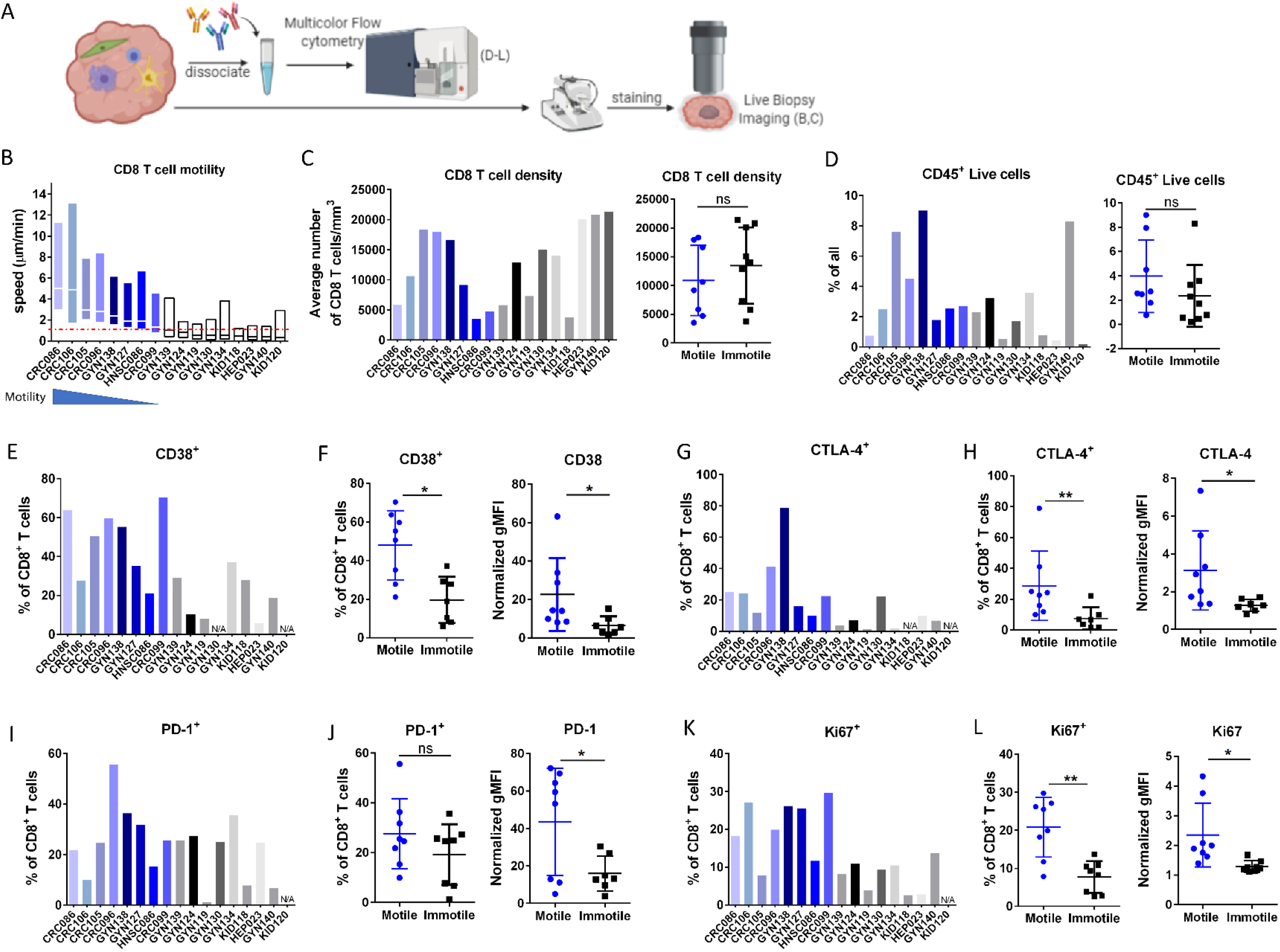
Linkage of T cell motility and exhaustion phenotypes in human tumors. (A) Diagram of a fresh tumor split for live biopsy imaging (panel B and C) and flow cytometry (panel D-L). (B) The speed of CD8 T cells across multiple tumor biopsies. Specimens with motile T cells were showed in blue bars; the immotiles (speed mean less than 1 μm/min) in black/grey. Red dashed line showed the cutoff. Each bar represents the CD8 T cell motility data pooled from at least three ROIs of one biopsy. Over 500 events were included for most of the biopsies. Min to Max is showed with line at mean. CRC: colorectal; HNSC: head and neck; GYN: Gynecologic; KID: Kidney; HEP: Hepatic. (C) Average CD8 T cell density as calculated by number of T cells per cubic millimeter pooled from multiple ROIs of a tumor. T cell density was plotted as the mean of CD8 T cell motility decreased (left). Tumor cases were also grouped based on T cell speed as motile or immotile (right). (D) Percentage of CD45^+^ live cells of the same tumor biopsies from panel A and B quantified by flow cytometry. (E-L) Flow cytometry of cells from the same tumor biopsies from panel A and B showed the percentage of CD38^+^, CTLA4^+^, PD-1^+^ and Ki67^+^ cells out of CD8 T cells. Percentage data plotted as both bar graphs and grouped into motile or immotile groups to compare. Normalized gMFI of these markers was also plotted for motile and immotile groups. Each dot represented percentage or MFI of a defined population from one biopsy._ * P<0.05, ** P<0.01 as determined by the Student’s t-test. N =17 (B – L).

We then turned to markers of exhaustion, guided by the mouse data (Fig 1), as a hypothesis. We gated samples for CD8 T cells and examined the frequency of cells expressing the exhaustion markers CD38 and CTLA-4, finding that these were upregulated in CD8 T cells in the motile group when compared to their expression in the stable group (Fig. 3E-H). PD-1 was also mildly upregulated in the motile group, however, was not as significant as CD38 and CTLA-4 (Fig. 3I, J).

Interestingly, when we derived a metric for macrophage maturation by ratioing the fraction of tumor associated macrophages (TAM) to their precursor classical monocytes (Mono), we found that increased values for that ratio also corresponded to increased motility (Supplemental Fig. 4E). This may represent a co-evolution of these two states, as is suggested by their co-distribution across tumor space (29).

We also sought to determine whether motility and exhaustion had other correlates and found that Ki67 expression in CD8 T cells followed the same trend (Fig. 3K, L). Though Ki67 is often used as a marker for proliferative capability, it also marks a population of recently generated terminal exhausted T cells that lose the ability to respond to additional stimulation (30, 31). From the tumor biopsies that we imaged, Ki67^hi^ CD8 T cells showed significantly higher expression levels of both PD-1 and CD38 than CD8 T cells from the same tumors with low expression of Ki67 (Supplemental Fig. 4F).

To further examine the link between T cell exhaustion and motility, we performed RNA-Seq of T cells sorted from a separate cohort of colorectal cancer patients, one of the most abundant indications among the cohort that has been analyzed by live biopsy as described in Figure 3. Through grouping the samples from this cohort based on CD38 upregulation—using CD38 as a representative marker for samples enriched for exhausted T cells — the top differentially upregulated genes were identified (Fig. 4A). *ENTPD1*, *LAG3*, *HAVCR2*, and *TOX2* were upregulated, supporting that these cells are preferentially exhausted whereas *TNF* was downregulated, indicating a reduced effector phenotype. Referencing gene ontology of the differentially expressed genes also showed that protein localization to microtubule and regulation of cell migration related pathways were both enriched in CD38 high group (Fig. 4B). These included genes that are associated with cytoskeletal rearrangement and cell polarization, e.g. *PDLIM4* (32), *AFAP1L2* (33), and *PLK4* (34), microtubule motor protein-encoding genes, *KIF20A* (35), *MYO7A* (36) and *MYO7B*, and a Rho-GTPase associated protein *CAV-1* (37) (Fig. 4A). We then plotted fold change for these promotility genes that revealed in our data by comparing ‘exhausted’ versus ‘non-exhausted’ cells based on the RNA-sequencing data of four recent publications on exhaustion and found they were all upregulated in exhausted T cells (Fig. 4C). Specifically, *MYO7A*, *AFAP1L2*, *PDLIM4* and *SKA3* were enriched in dysfunctional CD8 T cells from lung cancer or melanoma (38–40) and *MYO7A*, *AFAP1L2*, *PDLIM4*, *SKA3*, *KIF20A* and *PLK4* were upregulated in analyses comparing intratumoral PD-1^+^ versus PD-1^−^ CD8 T cells sorted from non-small-cell lung cancer samples(41).

**Figure 4.**
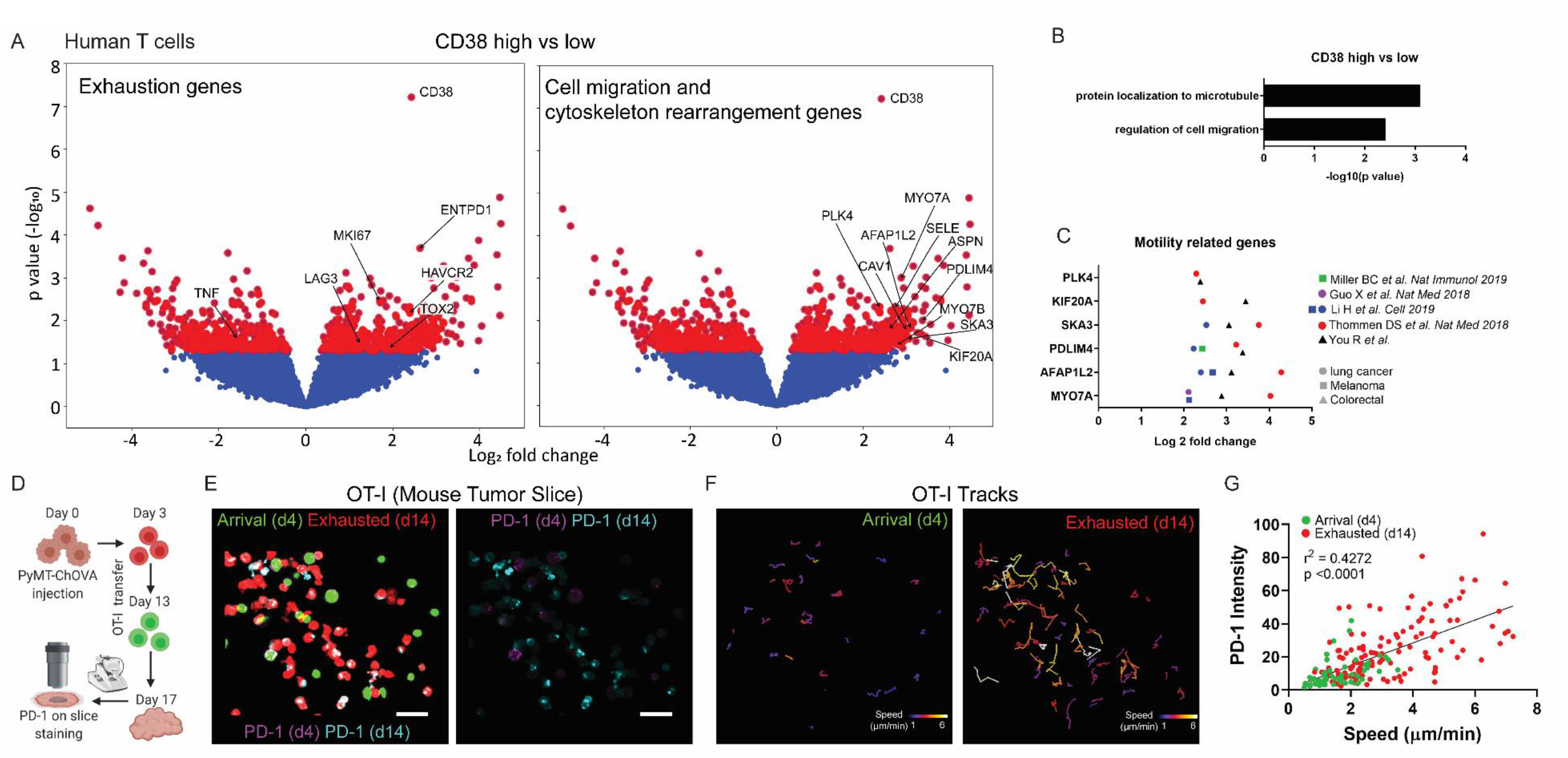
Exhausted T cells exhibited motile features. (A) Volcano plots of differentially expressed genes between CD38^hi^ (top 33^rd^ percentile of all samples for this gene) and CD38^low^ (bottom 33^rd^ percentile) in human T cells from 30 colorectal samples. T cell exhaustion genes are labeled in the left panel. Cell migration and cytoskeleton-related genes are labeled in the right panel. Red dots are above the cut off for p value = 0.05. (B) Gene ontology analysis of the upregulated genes in the CD38^hi^ samples compared to CD38^low^ samples, highlighting pathways related to cell migration. (C) Fold change of promotility genes in exhausted versus non-exhausted T cells from RNA-sequencing of melanoma or lung cancer samples in previous four publications. (D) Schematic diagram of the experimental design: RFP OT-I and GFP OT-1 were separately transferred into the same mice injected with PyMT-ChOVA cells in fat-pad at the time as indicated. Tumors were harvested for PD-1 on slice staining and imaging. (E) Representative images showing PD-1 staining on RFP OT-I cells (d14) and GFP OT-I cells (d4)_that resided in the same field. Scale bar = 30 μm. (F) Speed-based color-coded track displacement of GFP OT-1 (d4) on the left and RFP OT-1 (d14) on the right. Speed range is from 1 to 6 μm per minute. (E) Intensity of PD-1 staining on d4 (green dots) and d14 (red dots) OT-I cell was plotted against the mean speed of each cell. Data was pooled from three different ROIs and was representative of two independent experiments. P and r^2^ values were obtained by linear regression model.

To directly compare markers of exhaustion with cell motility on a cell-by-cell basis, we used the PyMT-ChOVA breast tumor model and slice imaging strategy (see Figure 1) to monitor the T cell motility and the expression of exhaustion markers simultaneously in slice (Fig. 4D). With on-slice staining of PD-1 in the live slices, we found that long resident T cells expressed higher PD-1 when compared to the recent arrivals (Fig. 4E). They also moved faster with a longer track compared to recent arrived effector T cells (Fig. 4F). When quantifying these parameters on individual cells from the same tumor volumes, d14 T cells exhibited higher PD-1 expression and speed, while d4 T cells expressed less PD-1 and were slower, with a positive overall correlation between T cell motility and exhaustion (Fig. 4G). Together, this supports that ‘exhausted’ cells, though defective for sensitivity and ability to mount cytokine responses, are nevertheless upregulated for genes associated with tissue surveillance.

In summary, we developed a two-photon microscopy-based imaging method of tumor slices from fresh biopsies taken from cancer patients that maintained immune dynamics and spatiotemporal information. This provides a systemic live imaging strategy of tumor biopsies focused on in situ endogenous T cell behavior in human cancer and provides a view of regional and patient-specific variations in cellular dynamics and further elaborates our understanding of a critical T cell state, namely ‘exhaustion’.

To the point of methods development, the biology observed using these imaging conditions and non-stimulatory antibodies highly resembles that observed in mouse lymph nodes and in situ (5–8, 11, 12, 26, 27), suggesting that we’ve broadly captured the durable biology of these tissues. Though the full-size immunoglobulins used in our methods were shown to be non-disturbing, Fc region-mediated effects, including cross-linking induced cell activation and antibody internalization, could be unavoidable in the tissue and in turn may change cell function. To improve it, using a monovalent Fab or nanobodies would be ideal. With their small size, tissue penetration could even be better. While all tissues and regions were studied identically in this method, in future and as possible through clinical testing, it may be useful to establish additional monitoring of oxygenation and other features during imaging and match them to metrics obtained for that tissue in the patient. For example, although we optimized our slice imaging method through the alteration of temperature and O_2_ perfusion—consistent with the established importance of these (42) and to match all mouse models from in situ to slice models—we still cannot exclude the possibility that immune behavior was not perfectly recapitulated in slice imaging because vascular circulation and chemokine milieu in addition to enervation and lymphatic flow were lacking (43).

Prior to the last decade, tumors have been mostly grouped based on their origins or mutations that drive uncontrolled growth. With the emergence of effective immunotherapies, tumors have been subtyped dependent on their immune phenotypes (44, 45). The abundance and composition of the immune infiltration is often used for the classification of immune subtypes of the cancer, e.g. immune rich or poor, lymphocyte enriched or depleted. More important is the sublocalization, for example understanding whether cells are ‘excluded’ from tumors (46). Prior to this study, static spatial immune infiltration patterns across cancer indications have been suggested as biomarkers for patients with solid tumors (45). With both spatial and dynamic quantification in the various compartments of tumors revealed through live biopsy, we can now further classify these previously ignored aspects of tumor heterogeneity.

Regardless of different histological tumor type and their genetic variations, th<u>e</u> combination of immune profiling by flow cytometry/RNA-Seq together with live biopsy imaging enabled us to group tumors according to T cell motility and link this to the aggregated state of the T cell population. In our study, the aggregated abundance of human exhausted T cells marked with Ki67^hi^ CTLA-4^hi^ PD-1^hi^ CD38^hi^ (17, 30) correlates well with the aggregated observation of fast motility within the same tumor sample. While multiple factors could contribute to the progression of T cell exhaustion, including varied levels of T cell priming in the lymph nodes, cellular and metabolite composition inside the tumor microenvironment, and desensitization to the existence of tumor antigens (47), our data suggested that T cells exhibited the exhausted phenotype gradually with a wide spectrum of exhaustion markers expressed at different levels. Correspondingly, though T cell speed within a sample varied, it was positively correlated to exhausted phenotype as a whole. Given the observation of an absolute shift in motility pattern during the establishment of exhaustion in mouse models (Figure 1 and 4) we believe that this relationship is likely to be at least partially cell intrinsic. This is further suggested by the upregulation of motility genes in T cells taken from patients with an abundance of exhausted cells. We cannot, however, exclude that some changes in motility during exhaustion result from differential priming in the lymph node, of some late-arriving cells. In addition, previous studies in islet tolerance suggest that PD-1 itself might block ‘stop’ signals generated by the TCR and so might be an additional component of what we observed here (48). Further analysis of single-cell sequencing data for exhausted T cells from patient samples, not currently available, may reinforce this as could in vitro experiments taken from patients. However, we consider it likely that this faster motility rate is a consequence of the combination of cell intrinsic and extrinsic features and may even represent the co-evolution of microenvironmental features along with the induction of exhaustion. Examples that regulate T motility include but are not limited to the chronic desensitization to persistent level of antigen (47), HLA expression level on tumor cells and APCs, and surface molecules that contribute to cellular interactions. With our live biopsy method established, it will be interesting to test if blocking TCR/HLA interaction could lead to the increase of motility or in contrast, if adding exogenous peptide or using anti-PD-1 blockade could arrest exhausted T cells in tumors. It will be even more intriguing to explore how other micro-regional factors, including the presence of specific cell types and soluble factors, such as chemokines, local oxygen levels, protease, and metabolites, regulate immune cell behavior. Additionally, how the micro-regional environment, like tumor or stroma density, myeloid cell type and abundance, and HLA expression level, affects T cell function, e.g. exhaustion, is to be determined. Due to the limited fluorophore options that are compatible with two photon and/or technical difficulty of the post-staining of the imaged slices that have been very well stabilized in the perfusion chamber with vet-bond and were hard to be removed intact, we did not perform any additional staining post live imaging to identify additional contributing factors that regulate local T cell motility and exhaustion .To understand all of these factors together, significantly more studies will be required, notably those that study spatial transcriptomics and/or proteomics.

Given that the composition and type of immune cells in the TME affect clinical outcomes (49), live biopsy developed in our study will provide a platform to test drugs, e.g. immune checkpoint inhibitors in tumors demonstrating these distinct T cell phenotypes. Following from the disseminated use and elaboration of these methods, live biopsy studies may act as mini-patients, to predict the efficacy of the same therapy in patients.

## Methods

### Patients and samples

Patients were enrolled in this study and provided written and informed consent to tissue collection under a University of California, San Francisco (UCSF) institutional review board (IRB)-approved protocol (UCSF Committee on Human Research (CHR) no. 13-12246). The clinical information of the patient samples used for imaging is summarized in Patient Information Table. The study enrollment period occurred from September 2017 to June 2018, and the sample size was determined by the availability of specimens throughout this period. The fresh biopsy samples were submerged in L15 medium and placed in a container of wet ice for transport to the lab within 5 hours.

### Human tissue slice staining and two photon imaging

Human tonsil or tumor biopsies were embedded in 2% low-melting agarose in PBS. Sections with 400 μm thickness were cut with Compresstome VF-200 (Precisionary Instruments Inc.) tissue slicer. Slices were stained with antibodies as mentioned including anti-CD8-Qdot605 (3B5), anti-CD14-Qdot705 (TüK4), anti-HLA-DR-Qdot705 (Tü36), anti-CD45-Qdot605 (HI30) and anti-Epcam-AF488 (9C4), for 1 hour in pre-oxygenated RPMI-1640 (phenol-red free) media at 37 °C with 5% CO_2_. Slices were washed in RPMI-1640 and attached to plastic coverslips using Vetbond (3M). Slices were kept at 37 °C in a heated-perfusion chamber with constant flow-over of RPMI-1640 (phenol red free) with carbogen (95% O_2_, 5% CO_2_) bubbled in throughout imaging. The MaiTai laser (Spectra Physics) was tuned to 780 nm for excitation of the Qdot fluorophores and second harmonic generation. The Chameleon laser (Coherent) excitation was tuned to 800 nm for excitation of AF488. Emitted light was detected using a 25X/1.2NA water lens (Zeiss) coupled to a six-channel detector array (custom; using Hamamatsu H9433MOD detectors), which were violet 417/50, blue 475/23, green 510/42, yellow 542/27, red 607/70, and far red 675/67. The microscope was controlled by the MicroManager software suite (28), and time-lapse videos were acquired at a frame rate of every 90 seconds or as indicated with 4-fold averaging at a z-resolution of 4 μm. Data analysis was performed using the Imaris software suite (Bitplane).

### Cell line and tumor injection

PyMT-ChOVA cell line was derived from PyMT-ChOVA spontaneous mammary tumor. Tumors were minced into small fragments and were cultured in DMEM plus 10% FCS with 1% penicillin–streptomycin– glutamine. Tissue fragments and debris were washed out with ice cold PBS and attached cells were kept in culture to confluency. Cells were then cultured for an additional 3 to 5 passages to generate PyMT-ChOVA cell line. For tumor cell injection, 2 × 10^5^ tumor cells were suspended in PBS and mixed 1:1 with Matrigel GFR (Corning) for a final injection volume of 50 μl. Cells were then injected in fat pad of CD2-dsRed mice anesthetized with isoflurane (Henry Schein).

### Two-photon intravital imaging of mouse tumor and analysis

Tumor-bearing mice were kept under anesthesia using isofluorane and mammary tumors were surgically exposed. Two-photon imaging was performed using a custom-built instrument equipped with 2 Ti-Sapphire lasers and 6 acquisition channels: laser 1, The MaiTai laser (Spectra Physics) was tuned to 800 nm (second harmonic generation, GFP, and AF647); The Chameleon laser (Coherent) was tuned to 980 nm (dsRed and mCherry). Time-lapse videos were acquired at a frame rate of every 90 seconds at a z-resolution of 3 μm. Data were analyzed using Imaris (Bitplane), including drift correction, video generation, cell surface detection, and generation of cell speed and track displacement data.

### Statistical Analysis

Statistical analyses were performed using GraphPad Prism software. Data are shown as mean ± s.d., calculated using prism. Specific statistical tests used were unpaired or paired Student’s *t*-test as indicated in the figure legend with p value described in detail. P and r^2^ values in Figure 4 were obtained by linear regression model.

**Supplemental Information** includes Supplemental figures, methods and videos can be found in the Supplementary material.

## Author Contributions

R.Y. and M.F.K designed experiments. R.Y. performed experiments unless specified. J.A., A.F. and A.E. assisted in two-photon microscopy experiments and imaging analysis. A.C. and G.R. participated in the design, preparation and analysis of experiments related to the flow cytometry of human samples. B.S processed and analyzed RNA-seq data. R.Y. and M.F.K wrote and revised the manuscript.

## Acknowledgements

We thank the Biological Imaging Development Center at the University of California San Francisco (UCSF) for help with microscopy data collection and instrumentation. We also thank the Parnassus Flow Cytometry Core for flow cytometry instrumentation, supported by grant no. P30DK063720 and the Institute for Human Genomics at UCSF for sequencing and bioinformatics support. Acquisition of certain human samples described in this study was through the collaboration with the Immunoprofiler Consortium at UCSF. This work was supported in part by the NIH/NCI grant no. R21CA196468 (M.F.K).

