## Supplementary Material for "Active Surveillance Characterizes Human Intratumoral T Cell Exhaustion"

**Supplemental Figures**

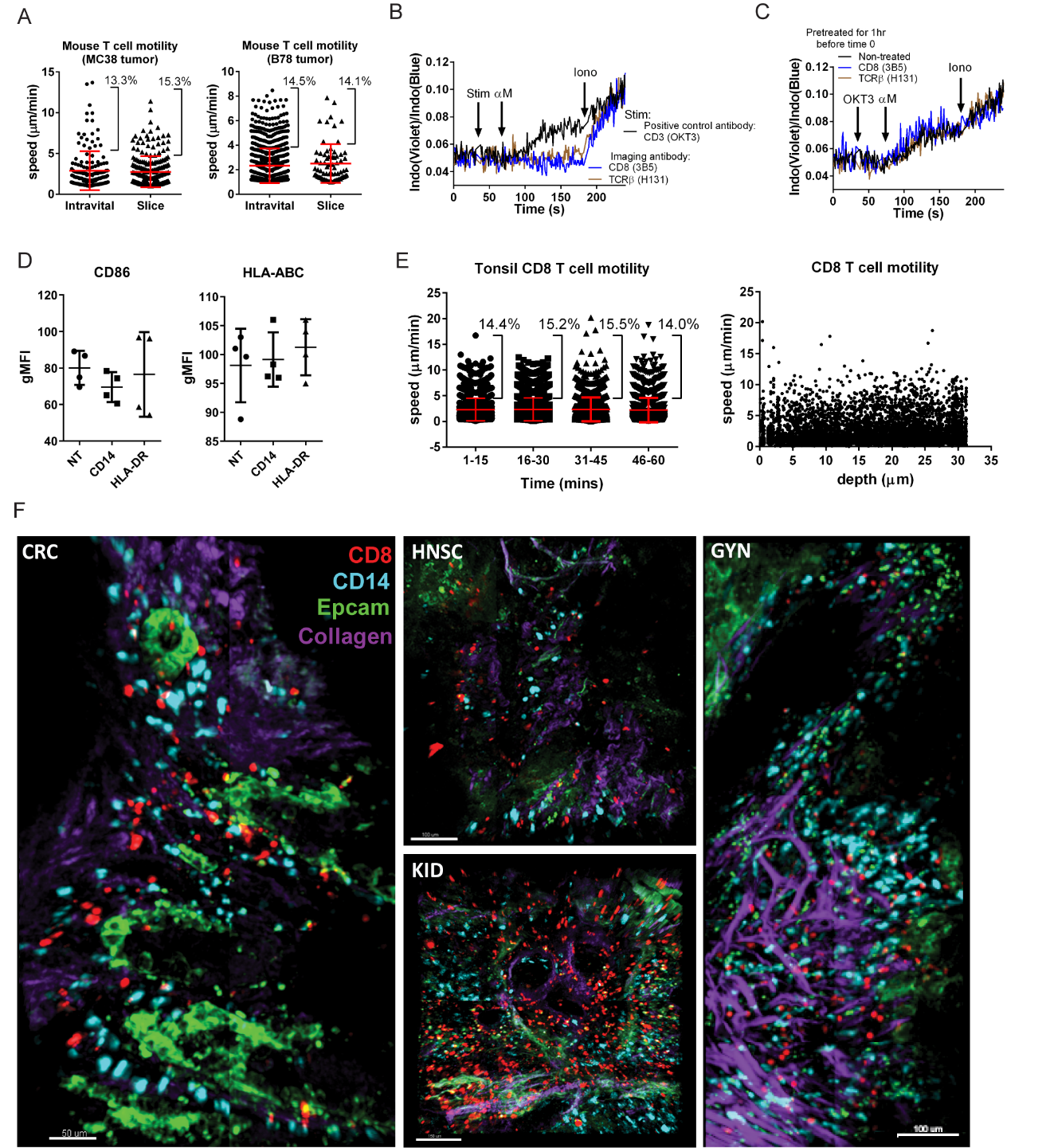

**Supplementary Figure 1. Establishment of live biopsy imaging** (related with Figure 1)**.** (A) Scatter plot showing the instantaneous speed of the individual CD8 T cell by intravital or slice imaging of the same tumor. The percentage of fast-moving T cells (Speed is over Mean+1 SD) was marked. (B) Calcium influx assay showed the activation of CD8 T cells in PBMC. Violet to blue fluorescence ratio indicates the calcium binding of Indo dye. Antibodies as indicated were added at 30 sec as stimulation reagents (stim) followed by anti-mouse IgG (αM) to cross link the antibodies. Ionomycin (Iono) were added at 200 sec as positive controls. (C) PBMC were pretreated with the antibodies as indicated followed by calcium influx assay. Anti-CD3 (OKT) antibody was used as a stimulatory antibody. (D) PBMC were pretreated with anti-CD14 of anti-HLA-DR antibody for 1 hour at 37 °C. The expression level of the indicated markers was quantified on myeloid cells (CD11b^+^CD15^-^CD3^-^CD19^-^ cells) in PBMCs by flow cytometry analysis. (E) Scatter plot showing the instantaneous speed of the individual CD8 T cell across 60 mins imaging time (left) and 30 μm imaging depth (right). Imaging depth was acquired based on the Z position of the specific T cell. Imaging time was divided into 15 mins-bins. (F) Representative images of colorectal (CRC), head and neck (HNSC), kidney (KID) and gynecological (GYN) tumor slice stained with anti-CD8-Qdot605, anti-CD14-Qdot705, and anti-Epcam-AF488. All images were acquired with two-photon microscopy with laser excitation tuned at 780 nm and 800 nm. Images were generated by Imaris software.

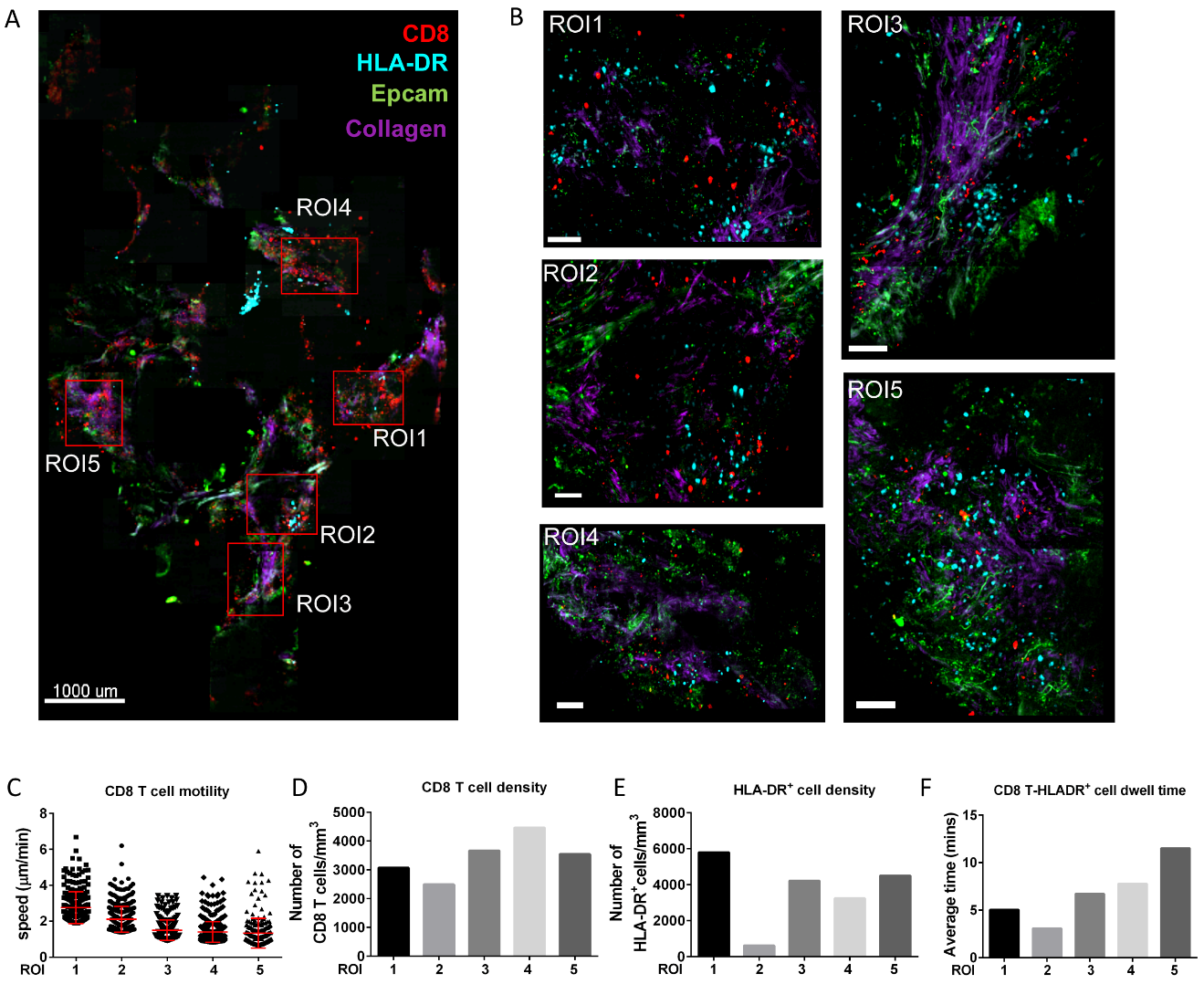

**Supplementary Figure 2.** **Diverse immune cell behavior in individual cancer biopsies.** (A) Representative image of human head and neck cancer slice stained with anti-CD8-Qdot605, anti-HLA-DR-Qdot705 and anti-EpCAM-AF488. (B) Five ROIs of panel A are showed. (C) The speed of top 10% fastest CD8 T cells from five ROIs was plotted with mean±SD. CD8 T cell (D) and HLA-DR^+^ cell density (E) was calculated by the number of cells per cubic millimeter. (F) The dwell time between CD8 T cell and HLA-DR^+^ cell was calculated by the average duration of two types of cells in contact with each other. All images were acquired with two-photon microscopy with laser excitation tuned at 780 nm and 800 nm. Images were analyzed by Imaris software. Dwell time was quantified with a MATLAB extension to Imaris software.

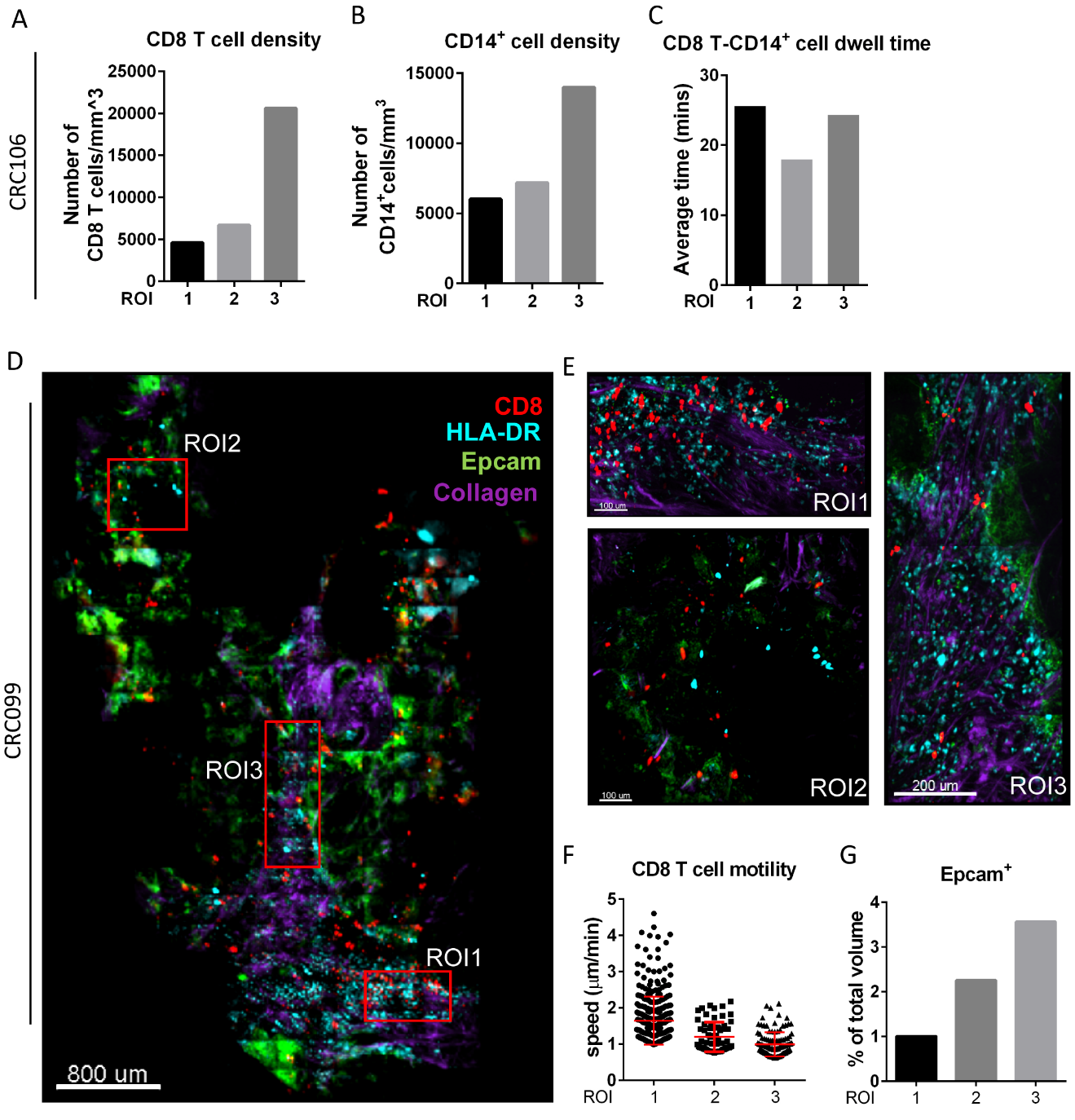

**Supplementary Figure 3. Live biopsy of colorectal cancers** (related with Figure 3)**.** CD8 T cell (A) and HLA-DR^+^ (B) cell density is calculated by the number of cells per cubic millimeter in three ROIs of the same colorectal tumor in Figure 3. (C) The dwell time between CD8 T cell and CD14^+^ cell is calculated by the average duration of two types of cells in contact with each other. (D) Overall view and (E) three ROIs of another colorectal cancer slice stained with antibodies anti-CD8-Qdot605, anti-HLA-DR-Qdot705, and anti-Epcam-AF488. CD8 T cell motility (F) and density of Epcam^+^ volume (G) of the tumor in panel D and E are quantified by Imaris software. All images were acquired with two-photon microscopy with laser excitation tuned at 780 nm and 800 nm. Images were analyzed by Imaris software.

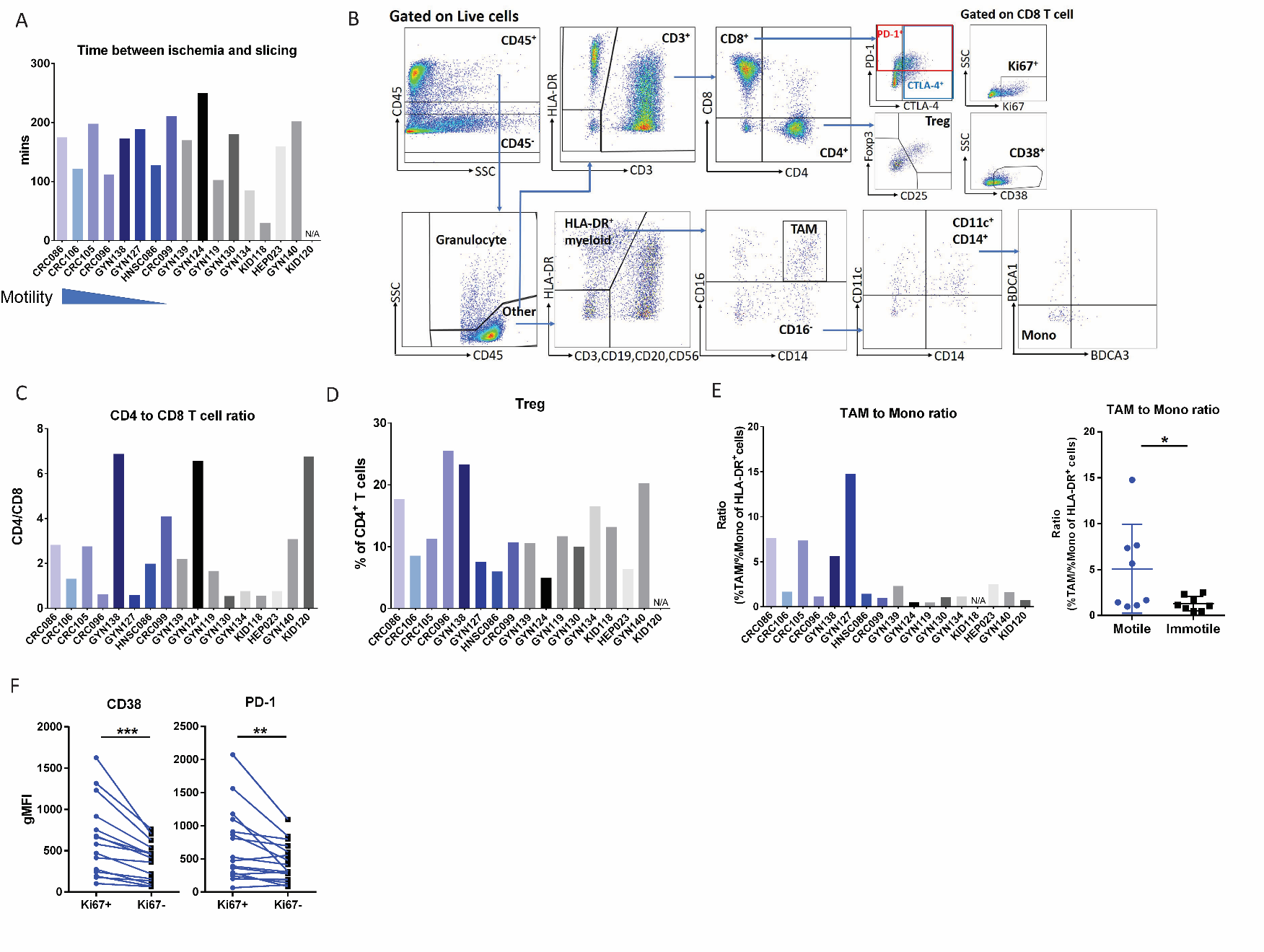

**Supplementary Figure 4.** **Immune profiling of cancer biopsies** **that were imaged with live biopsy** (related with Figure 4). (A) Tissue storage time for tumors used in live biopsy imaging. Time was plotted for each specimen. (B) Gating strategy of human tumor T cell and myeloid population. (C) The ratio of the abundance of CD4 to CD8 T cell and (D) regulatory T cells (Tregs) were plotted as the mean of CD8 T cell motility decreased. Blue bars represented specimens with motile T cells; black/grey bars represented those with immotile T cells. (E) The ratio of the percentage of tumor associated macrophages (TAM) of HLA-DR^+^ cells to the percentage of classical monocytes (Mono) of HLA-DR^+^ cells was plotted as the mean CD8 T cell motility decreased and also plotted when all the samples were grouped into motile and immotile groups. (F) Geometric MFI of CD38 and PD-1 expressed in Ki67^+^ versus Ki67^-^ CD8 T cells from each sample. * P<0.05, ** P<0.01, *** P<0.001 as determined by the Paired Student’s t-test. N =17 samples.

**Supplementary Video 1.**

Live biopsy of a tonsil slice showed that T cells (red) moved freely (yellow arrows) or dwelled with APCs (cyan) (white arrow).

**Supplementary Video 2.**

Live biopsy of a colorectal cancer slice showed that 1) overall view, 2) zoomed in a tumor sparse area where T cells (red) migrated freely, and 3) zoomed in an area where T cells dwelled with tumor cells (green) and APCs (Cyan).

**Supplemental Materials and Methods**

**Mouse strains.** All mice were housed and bred at University of California, San Francisco (UCSF) and were maintained under specific pathogen-free conditions in accordance with the regulatory standards of the National Institutes of Health (NIH) and American Association of Laboratory Animal Care standards and that were consistent with the UCSF Institution of Animal Care and Use Committee (IACUC approval no. AN170208-01B). PyMT-ChOVA (C57BL/6) transgenic mice were generated as previously described ^1^. CD2-dsRed mice were obtained as described previously (MGI# 5296821) ^2^. Ubiquitin-GFP mice were obtained from The Jackson Laboratory. OT-I mice (The Jackson Laboratory) were bred to CD2-dsRed and Ubiquitin-GFP mice to generate mice used for OT-I transfer experiment.

**Calcium influx assay.** Jurkat cells or PBMCs were loaded with Indo-1 calcium indicator (Thermo Fisher Scientific) according to manufacturer’s instruction. Cells were warmed to 37°C for 5 mins and applied to flow cytometer to record baseline of calcium level. Antibodies were added as described during the process and calcium influx was monitored.

**OT-I adoptive transfer for imaging and flow cytometry analysis.** 5 x 10^6^ RFP OT-I cells or GFP OT-I isolated from LN using CD8 EasySep enrichment kit (STEMCELL Technologies) were transferred into PyMT-ChOVA-bearing mice at 14 days or 4 days respectively before imaging and flow cytometry experiments. Tumor sections with 400 μm thickness were cut with Compresstome VF-200 (Precisionary Instruments Inc.) tissue slicer and stained with PD-1 Alexa Fluor 647 antibody (clone 29F.1A12, 2.5 µg/ml, BioLegend 135230) for 1 hour at 37 °C. Slices were imaged as described in the human tissue slice imaging session. PD-1 intensity was generated by Mean Fluorescent Intensity of AF-647 on RFP or GFP OT-I T cells using Imaris Software. The rest of the same tumors were minced, and incubated in digestion buffer (100 U/ml collagenase type I (Roche), 500 U/ml collagenase type IV (Roche), and 200μg/ml DNAse I (Roche) in RPMI-1640 (GIBCO)) for 30 mins on a shaker at 37 °C. Cells were then washed with PBS plus 2% FCS and filtered through a 100 μm cell strainer before staining for flow cytometry.

**Human tissue digestion**. The fresh biopsies were minced with surgical scissors and transferred to a GentleMACs C Tube (Miltenyi Biotec) with 100 μg/mL Liberase TL (Roche) and 200 μg/mL DNase I (Roche) at 3 ml per gram of tissue. C tubes were incubated in the GentleMACS octo dissociator (Miltenyi Biotec) with heaters, following the manufacturers dissociation protocol (Miltenyi Biotec Tumor Dissociation Kit). 10 mL of sort buffer (PBS + 2% fetal calf serum (FCS) + 2 mM EDTA) was added to samples and filtered through a 100 μm filter and spun down, and red blood cells were lysed with 175 mM ammonium chloride. Samples were then filtered through a 70 μm filter, spun down, and resuspended for staining ^3^.

**Antibodies against human antigens**. The following antibodies were used: TCR clone C305 (Sigma-Aldrich 05-919), TCR Vβ13.1 clone H131 (BioLegend 362403), TCR α/β clome IP26 (BioLegend 306711), CD45 clone Hl30 (eBioscience 47-0459-42 and Thermo Fisher Scientific Q10051), CD3e clone OKT3 (eBioscience 46-0037-42), clone UCHT1 (BioLegend 300422) and clone HIT3a (Thermo Fisher Scientific 12-0039-41), HLA-DR clone L243 (eBioscience 48-9952-42) and clone TÜ36 (Thermo Fisher Scientific Q22159), CD56 clone CMSSB (eBioscience 46-0567-42), CD56 clone HCD65 (BioLegend 318304), CD19 clone H1B19 (eBioscience 46-0198-42 and 56-0199-42), CD14 clone M5E2 (BioLegend 301836), CD16 clone 3G8 (BioLegend 302040), CD11c clone 3.9 (eBioscience 56-0116-42), CD85g clone 17G10.2 (eBioscience 12-5179-42), CD4 clone RPA-T4 (BioLegend 300550), CD8 clone RPA-T8 (BioLegend 301040) and clone 3B5 (Q10009), CD25 clone BC96 (BioLegend 302612), CD25 clone 2A3 (eBioscience 17-0259-42), FOXP3 clone PCH101 (eBioscience 77-5776-40), PD-1 clone EH-12 (BioLegend329930), CTLA-4 clone BNI3 (BioLegend 369606), γδ T cell receptor (TCR) clone B1.1 (BioLegend 331212) and CD326 (EpCAM) clone 9C4 (BioLegend 324210). CD38 clone HIT2 (BD Biosciences 564980).

**Antibodies against mouse antigens**. The following antibodies were purchased from Biolegend and used: CD90.2 AF700 clone 30-H12, CD11b BV785 clone M1/70, CD279/PD-1 BV605 clone 29F.1A12, and IFNγ APC clone XMG1.2. Ki-67 PE-eFluor 610 clone SolA15 was from ThermoFisher Scientific. CD38 BV711 clone 90 was from BD Biosciences. TOX APC clone REA473 was from Miltenyi Biotec.

**Flow Cytometry.** For surface staining, cells were incubated with Fc receptor–blocking solution (Human TruStain FcX, BioLegend 422301 or anti-Fc receptor antibody (clone 2.4G2) for mouse samples) and stained with antibodies in PBS + 2% FCS + 2 mM EDTA for 30 min on ice. Viability was assessed by staining with Zombie Aqua fixable viability dye (BioLegend 423102) or Zombie NIR fixable viability dye (BioLegend 423106). All human samples were fixed with BD Cytofix before analysis, following the manufacturer’s protocol. For intracellular staining, samples were first stained for surface markers, as described above, and then fixed with fixation/permeabilization solution and washed with permeabilization buffer, per the manufacturer’s suggested protocol (eBioscience 00-5523-00). Intracellular staining was undertaken in the presence of 2% FCS and an Fc receptor–blocking solution. Flow cytometry was performed on a BD Fortessa or an Aria Fusion flow cytometer. Analysis of flow cytometry data was done using FlowJo (Treestar) software.

**RNA sequencing analysis of human tumor samples**. With each colorectal tumor sample, 2,000 to 10,000 live T cells were directly sorted into lysis buffer by fluorescence-activated cell sorting (FACS) to isolate RNA. RNA was isolated using the Dynabeads mRNA DIRECT purification kit (ThermoFisher Scientific 61011), per the manufacturer’s suggested protocol. Isolated RNA was converted into amplified cDNA using the Ovation RNA-Seq System V2 kit (NuGen 7102-A01), and cDNA was converted into RNA-seq libraries using the Nextera XT DNA Library Prep kit (Illumina FC-131-2001), following the manufacturer’s suggested protocol. RNA-seq library quality was checked using a Bioanalyzer HS DNA chip (Agilent) and pooled. Pooled libraries were then sequenced via a single-read 50-bp MiSeq (Illumina) run, and libraries containing > 10% of reads aligned to coding regions and > 1,000 unique reads in the total library were selected for further sequencing. RNA-seq libraries that met the previous quality control criteria were pooled and submitted to the UCSF Center for Advance Technology for paired-end, 100-bp (PE100) sequencing on the HiSeq4000 (Illumina). The generated fastq files were checked for the quality using fastq to trim nextera adapters and poly-G artifacts arising from low insert size fragments. The Burrows–Wheeler Aligner was used to align sequenced RNA-seq library reads against a reference of ribosomal RNA (rRNA) and mitochondrial encoded 12s and 16s ribosomal RNA to deplete the dataset of reads arising from these sequences. The remaining reads were then aligned to the human reference genome build GRCh38 (annotated with the Ensembl GrCh38.85 gtf) using the software package STAR. Gene expression was estimated from aligned BAMs using the software package RSEM. CD38 high samples were the samples in the top 33^rd^ percentile of all samples based on the CD38 transcript per million (TPM) while low expression samples were in the bottom 33^rd^ percentile of all samples. Identification of differentially expressed genes (DEGs) was performed comparing high and low populations with limma package ^4^ and voom ^5^, a function of limma that modifies RNA-Seq data for use with limma.
